# Balanced polymorphisms in gamete-binding genes are not associated with human infertility

**DOI:** 10.1101/2024.10.11.617896

**Authors:** M. W. Hart, B. Crespi, K. Seethram

## Abstract

Genes expressed in gametes that mediate sperm interaction with the mammalian egg are of considerable interest to evolutionary biologists because the evolution of such genes can account for variation in reproductive compatibility between mates and reproductive isolation between species. The human orthologs of such genes are also potential targets for both contraception and treatment of infertility. One gene system of particular interest is the sperm-binding genes of the inner egg coat or zona pellucida (*Zp2, Zp3*) and their cognate protein in the mouse sperm acrosome (*Zp3r*). Previous population genetic analyses in humans pointed toward three balanced polymorphisms (one in each gene *ZP2, ZP3*, and *ZP3R*) as potential targets of some form of balancing selection in the evolution of human fertility. We tested that association using genetic analysis of couples seeking fertility assistance, but we could not reject the null hypothesis of no association between balanced polymorphisms and infertility. Our study was based on a small sample of couples, but the data were sound: the allele frequencies at those three balanced polymorphisms were not different from random expectation in that clinical sample. If an effect of those allele frequencies on infertility exists it is probably small. Our study was based in part on an old error in gene annotation that was only recently discovered (after the start of participant recruitment for this genetic analysis), and this error may account for our results, which argue against a role for balancing selection on those three genes in humans.

## Introduction

Gamete interactions at fertilization affect reproductive success between mates and reproductive isolation between populations or species (Carlisle & Swanson, 2021). Understanding those interactions in humans and other mammals has important implications for the development of contraceptives and fertility treatments (Meeusen et al., 2007; Hirohashi et al., 2008; Huang et al., 2014). Previous experimental studies and evolutionary analyses in both mice and humans suggested that interactions between a sperm protein in the acrosome (ZP3R) and two proteins in the egg vitelline envelope or zona pellucida (ZP2, ZP3) affect binding between gametes (Buffone et al., 2008; Avella et al., 2013). The evolution of such systems of genes is commonly influenced by sexual selection acting on differences between males and females in optimal values for traits such as the frequency or quality of interactions between sperm and eggs (Vacquier, 1998; Gavrilets, 2000; van Doorn et al., 2001; Swanson & Vacquier, 2002; Turner & Hoekstra, 2008; Okamoto, 2016; Evans & Lymbery, 2020; Weber & Fisher, 2023). One common outcome of such selection is the evolution of differences in compatibility between some combinations of paternal and maternal genotypes (Palumbi, 1999; Levitan, 2012; Hart et al., 2014; Levitan et al., 2019).

Evidence for such selection includes linkage disequilibrium (LD): unexpected high frequencies of genotype combinations among unlinked genes that encode interacting fertilization proteins (Tomaiuolo & Levitan, 2010; Stapper et al., 2015). Such combinations are expected to be broken up by independent assortment and recombination during meiosis each generation, so the maintenance of such combinations over multiple generations requires some deviation from neutral population genetic expectations (Charlesworth et al., 1997). Linkage disequilibrium between unlinked fertilization genes might be maintained by balancing selection favouring some genotype combinations over others due to differences among pairs of mates in gamete compatibility (Kekäläinen, 2021). Balancing selection is of general interest to evolutionary ecologists because it can account for the maintenance of otherwise unexplained genetic variation (Mitchell-Olds et al., 2007; but see Zietsch, 2024).

Two previous analyses suggested that a human homolog of mouse *ZP3R* may coevolve under selection with human *ZP2* and *ZP3*. Rohlfs et al. (2010) showed that single-nucleotide polymorphisms (SNPs) in the human gene they identified as *ZP3R* on chromosome 1 were in LD with SNPs in human *ZP3* on chromosome 7 in a large and diverse nonclinical subpopulation. Hart et al. (2018) followed up that discovery by quantifying LD and other evidence of selection acting on one balanced polymorphism (with high minor-allele frequency) in each of those genes—plus a third balanced polymorphism in *ZP2* on chromosome 16—in the 1000 Genomes data, including an association between measures of fertility and genotypes at two of those loci in a small founder subpopulation. Authors of those two studies proposed that those population genetic patterns could arise from selection on interactions between those protein pairs during sperm binding to the egg coat.

If the fertility effects of those protein interactions are strong enough to influence population genetic variation in nonclinical population samples from fertile individuals and families, then evidence for that effect is expected to be detectable in clinical samples from couples seeking treatment for infertility. In this study we tested that prediction among couples seeking treatment at a large urban fertility clinic. We sampled genetic variation at one polymorphic site in each gene that is putatively under selection, and used two tests of the prediction that frequencies of male-female genotype combinations are shaped by selection for fertility. We did not find evidence that genotype combinations in couples are associated with infertility. We discuss the broader context for that discovery including recent analyses of human *ZP3R* that point to an error in the conceptualization of our genotyping study.

## Methods

We used qPCR to genotype 92 individuals at three well-studied human single-nucleotide polymorphisms (SNPs) in *ZP3R* (rs4844573), *ZP2* (rs2057720), and *ZP3* (rs2906999) that were previously observed to be in LD and associated with fertility variation in non-clinical population samples. We recruited couples seeking fertility assistance at the Pacific Centre for Reproductive Medicine (PCRM, Burnaby, British Columbia). Because we were interested in a hypothesis about compatibility between eggs and sperm we limited recruitment to couples consisting of a female and a male partner. Informed consent to participate in the study was collected from each individual in each couple by PCRM staff. Methods for recruitment of participants, collection of samples, and confidential handling of participant data were approved by the Simon Fraser University Office of Research Ethics (H20-00273-A007). A total of 46 couples agreed to participate, including 37 couples in the experimental arm of the study (couples unable to achieve a pregnancy, possibly including some who experienced infertility due to sperm–egg incompatibilities involving ZP3R, ZP2, and ZP3), and 9 couples in a control arm of the study (couples who were able to achieve a pregnancy, but unable to carry the pregnancy to full term). In the original conceptualization of the study, we hoped to do a direct comparison of genotype combinations between those two groups of couples (from the same subpopulation of couples in Burnaby and the surrounding urban region seeking treatment for infertility), but the small size of the control arm limited our analyses to tests of predictions based only on the experimental arm of the study.

Each individual submitted a sample of cheek epithelial cells that was self-collected and returned to PCRM for genotyping at Simon Fraser University. Cell samples were collected using OCR-100 sample collection kits (DNA Genotek). Genomic DNA was extracted from each sample using prepIT-L2P DNA extraction reagent (DNA Genotek). Each DNA pellet was washed with ethanol and dissolved in water. Genomic DNA concentration was checked on a NanoDrop 2000c spectrophotometer and diluted to ∼15 ng/µl. We genotyped each sample using two off-the-shelf qPCR assay probes for *ZP3R* (ThermoFisher catalog number 4351379; assay ID C_32309470_20; SNP ID rs4844573) and *ZP2* (catalog number 4351379; assay ID C_2607578_1; SNP ID rs2075520).

To assay the third polymorphism for *ZP3* we used a custom assay kit manufactured by ThermoFisher to mask another SNP (rs148976073) at the genomic location (chr7:76440493) next to the balanced polymorphism rs2906999 (chr7:76440494) we intended to genotype. We tested that custom assay kit against 10 samples of individuals from the CHS subpopulation in the 1000 Genomes biobank at the Coriell Institute that have known rs2906999 genotypes including three CC homozygotes (HG00404, 422, 428), three TT homozygotes (HG00407, 421, 427), and four CT heterozygotes (HG00403, 406, 533, 534).

We used those three assay kits in 10-µl qPCR assays containing 0.6 µl genomic DNA, 5 µl TaqMan Genotyping Master Mix (ThermoFisher), 0.5 µl of 20x assay probe, and 3.9 µl water. We detected VIC and FAM fluorescence (indicating the presence of one or both alleles at each SNP) for each sample and each assay probe on a LightCycler96 instrument. We used two-step (*ZP3R, ZP2*) or three-step (*ZP3*) thermal cycling for 55– 80 cycles to obtain fluorescence signals and analyzed the fluorescence data using LightCycler96v1.1 software.

We used two methods to test the prediction that genotype incompatibilities between mates might cause those 37 couples to seek fertility assistance at PCRM. First, we calculated the frequencies of the two alleles at each SNP in the sample of 184 alleles (2 alleles for 92 individuals in 46 couples) and used those allele frequencies to calculate the expected frequency of genotype combinations among the 37 couples in the experimental arm of the study if those combinations resulted from a random draw of alleles and genotypes from the GVRD population. We used a goodness-of-fit test to compare the expected frequencies of genotype combinations (from the allele frequencies in the population) to the observed frequencies of families with each male *ZP3R* genotype expressed in sperm and each female *ZP2* and *ZP3* genotype expressed in the egg coat (in the sample of 37 couples from the experimental arm of the study). We compared the resulting test statistic to the critical value of the *χ*^2^ distribution with 16 degrees of freedom: each family must have 1 of 3 male genotypes (for *ZP3R*) and 1 of 9 different two-locus female genotypes (for *ZP2* and *ZP3*), giving (3-1)*(9-1) = 16.

Second, we carried out the same comparison between expected frequencies of genotype combinations (from the population allele frequencies) and observed frequencies of female *ZP3R* genotypes and male *ZP2 & ZP3* genotypes. This comparison provides a helpful control because the *ZP3R* gene is not expected to be expressed in the female member of each couple, and the *ZP2 & ZP3* genes are not expected to be expressed in the male member of each couple. Because those genotypes (her sperm genes, his egg coat genes) are not expressed in either individual, the genotype combinations are not expected to cause those couples to experience infertility or seek fertility assistance at PCRM.

Comparing the statistical results from the first analysis to the second analysis provides a within-sample control for non-random variation in the frequencies of male-female genotype combinations that is not caused by gamete incompatibility. To our knowledge, this is a novel approach to analyzing the potential influence of interacting genes with sex-specific expression on the fertility of male-female pairs.

## Results

We obtained high-quality genotype data for all 92 individuals at all three SNPs. An example of the genotype-calling output for one locus (rs4844573 in *ZP3R*) is shown in Figure 1.

**Figure 1.**
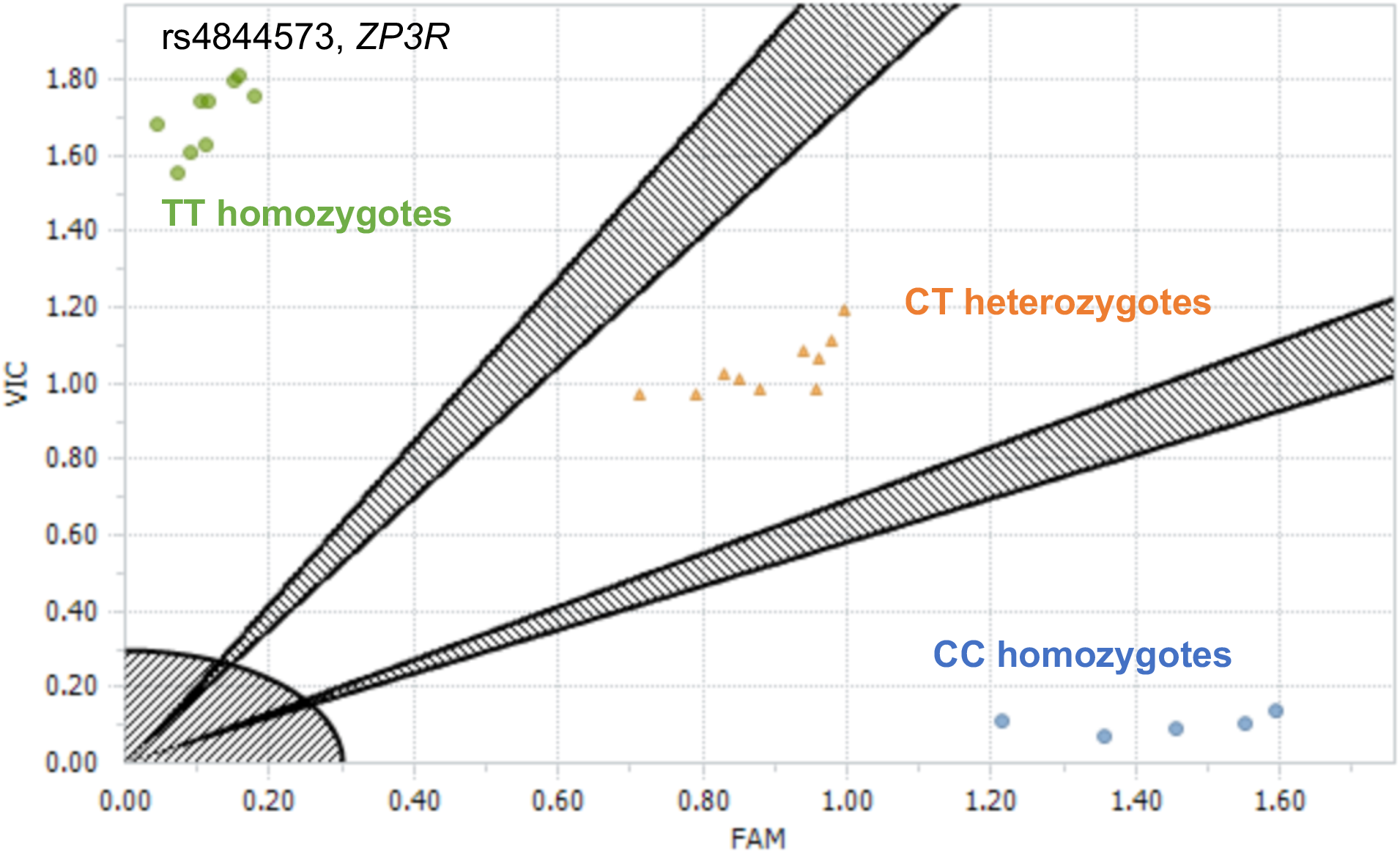
Example qPCR genotyping results based on relative fluorescence of two probes and fluorescent dyes (FAM, VIC) each associated with one nucleotide allele (C, T) at a single-nucleotide polymorphism (rs4844573, *ZP3R*). Clusters of individuals for which relatively high fluorescence was detected for only one of the two dyes share the same homogyzous CC (blue circles) or TT (green circles) genotype; individuals for which both fluorescent signals were detected share the heterozygous CT genotype (orange triangles).

At each SNP, the more common of the two nucleotide alleles occurred at a frequency of about 0.6 (Table 1), and the less common or minor allele occurred at a relatively high frequency (0.34–0.40). This high minor allele frequency is characteristic of those three SNPs across all human subpopulations.

**Table 1.** Counts of genotypes and alleles (and allele frequencies) for three balanced human polymorphisms in genes that mediate mammalian fertilization.

| Locus | Genotype count |  | Allele count |  | Allele frequency |
| --- | --- | --- | --- | --- | --- |
| rs2075520 (*ZP2*) | GG | 45 | G | 122 | 0.66 |
|  | TT | 15 | T | 62 | 0.34 |
|  | GT | 32 |  |  |  |
| rs2906999 (*ZP3*) | CC | 30 | C | 110 | 0.60 |
|  | TT | 12 | T | 74 | 0.40 |
|  | CT | 50 |  |  |  |
| rs4844573 (*ZP3R*) | TT | 39 | T | 119 | 0.65 |
|  | CC | 12 | C | 65 | 0.35 |
|  | CT | 41 |  |  |  |

We used those allele frequencies to predict the frequency of genotype combinations in a sample of 37 couples if those combinations came from a random draw of alleles from the sampled population (Table 2). Because our sample of couples was small, all expected frequencies of those genotype combinations were low and ranged from about 0.08 for the least common genotype combinations (e.g., the CC/TT/TT genotype combination composed of homozygotes for each of the three minor alleles) to about 3.6 couples (e.g., the CT/GT/CT genotype combination composed of heterozygotes for each of those balanced polymorphisms).

**Table 2.** Expected genotype counts among 37 sampled couples (from Table 1).

|  |  |  | **ZP3R** |  |  |  |  |  |  |  |  |
| --- | --- | --- | --- | --- | --- | --- | --- | --- | --- | --- | --- |
|  |  |  | *Thr* CC |  |  | CT |  |  | *Ile* TT |  |  |
| **ZP3** |  |  | *Pro* CC | CT | *Ser* TT | CC | CT | TT | CC | CT | TT |
| **ZP2** | *Gly* | GG | 0.71 | 0.95 | **0.32** | 2.64 | 3.52 | 1.17 | 2.45 | 3.27 | 1.09 |
|  |  | GT | 0.73 | 0.98 | 0.33 | 2.72 | 3.63 | 1.21 | 2.53 | 3.37 | 1.12 |
|  | *Val* | TT | 0.19 | 0.25 | 0.08 | 0.70 | 0.93 | 0.31 | 0.65 | 0.87 | 0.29 |

For the genotype combinations with relatively high expected frequencies, the interpretation of that expectation is straightforward: among 37 sampled couples, we expect to observe 3 or 4 couples with that most-common CT/GT/CT genotype combination. For the genotype combinations with relatively low expected frequencies (e.g., *f* = 0.08), one interpretation is that across many repeated samples of 37 couples from our study population, only about 1 in 12 of those repeated samples would be expected to include a couple with that rare homozygote CC/TT/TT genotype combination composed of only the less-common allele at all three loci.

The observed genotype combinations for the three genes that are expressed in sperm (male *ZP3R* genotype) and in eggs (female *ZP2*-*ZP3* genotype) were slightly different from those predicted frequencies (Table 3). The most common observed genotype combinations among the 37 couples in the experimental arm included some of the most common expected genotype combinations, and we observed few couples with genotype combinations that had low expected frequencies (consisting of a larger number of less common alleles). The goodness-of-fit test (*χ*^2^ = 28.90, *df* = 16) was marginally significant (*p* = 0.025).

**Table 3.** Observed genotype counts among 37 sampled couples for expressed genes.

|  |  |  | **ZP3R** |  |  |  |  |  |  |  |  |
| --- | --- | --- | --- | --- | --- | --- | --- | --- | --- | --- | --- |
|  |  |  | Males |  |  |  |  |  |  |  |  |
|  |  |  | *Thr* CC |  |  | CT |  |  | *Ile* TT |  |  |
| Females | **ZP3** |  | *Pro* CC | CT | *Ser* TT | CC | CT | TT | CC | CT | TT |
| **ZP2** | *Gly* | GG | 0 | 1 | **2** | 3 | 5 | 1 | 4 | 4 | 0 |
|  |  | GT | 0 | 0 | 0 | 3 | 2 | 0 | 0 | 6 | 0 |
|  | *Val* | TT | 0 | 1 | 0 | 1 | 1 | 1 | 0 | 1 | 1 |

That *p* value could be interpreted to mean that the 37 infertile couples in that sample show a significant difference from expected genotype frequencies that may be caused by selection on those genotype combinations during sperm-egg interactions and might be the cause of infertility. However, that conclusion is not very strong. The sample of infertile families is small (37) relative to the number of possible genotype combinations (27 different family combinations from 3 different male genotypes x 9 different two-locus female genotypes), so all of the observed and expected frequencies are low. These low values make the goodness-of-fit test sensitive to stochastic effects. For example, about one-third of that large *χ*^2^ value is associated with the 2 families with the CC/GG/TT genotype (bold value in Table 3) that is predicted to be rare (0.32 out of 37 families or about 1%; bold value in Table 2).

A second reason to treat that result with caution is that we observed a similar difference between observed and expected genotype combinations in the analysis of genotypes that are not expressed in gametes (female copy of the sperm gene, male copies of the egg coat genes) and cannot contribute directly to infertility (Table 4).

**Table 4.**
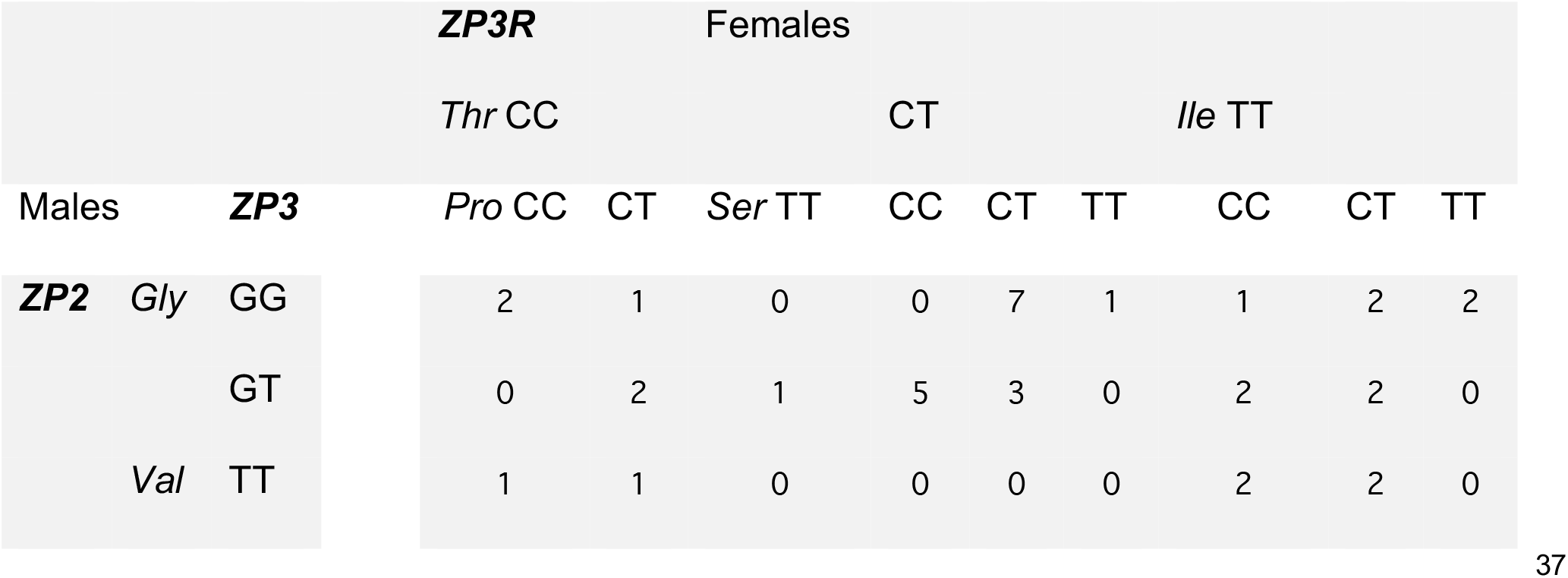
Observed genotype counts among 37 sampled couples for nonexpressed genes.

Like the analysis of the genotypes expressed in gametes, the goodness-of-fit test for this non-functional combination of genotypes was significant (*χ*^2^ = 31.41, *df* = 16, *p* = 0.011). Because those genotype combinations are not expressed in those 37 couples seeking fertility assistance, the significant test result cannot be interpreted as evidence that those genotype combinations contribute to infertility. The significant statistical result is most likely to be explained by small sample sizes and the sensitivity of the goodness-of-fit test to stochastic variation. That interpretation applies to both analyses of the genotypes expressed in gametes (Table 3) and not expressed in gametes (Table 4), and suggests that molecular interactions among these *ZP3R, ZP2*, and *ZP3* alleles at fertilization are probably not contributing to infertility of these couples.

## Discussion

Given the previous evidence for functional interactions among these genes in mice and coevolution among them in humans, and the evidence for an effect of two of these genes on fertility in a non-clinical population sample of humans, how can we account for these new results?

First, the genetic evidence for an effect of *ZP3R, ZP2*, and *ZP3* alleles on infertility is expected to be relatively weak in this sample because there may be several causes for the inability to achieve a pregnancy among the 37 couples in the experimental arm of the study, possibly including some who experience infertility due to genotype incompatibilities during sperm–egg binding. A much larger sample might be needed to detect that specific effect in a heterogenous sample of couples with a similar fertility phenotype but different physiological or molecular mechanisms underlying their infertility. However, the results from our study do not seem to point to any specific genotype combination that might be a possible source of infertility among some couples in this small sample

In a larger sample of Hutterite families (see Kosova et al., 2012), we previously noted a weak but significant association between the maternal *ZP2* G allele (rs2075520) and smaller family sizes (Hart et al, 2018). In this new sample of infertile couples, we might have expected to see (for example) a relatively high frequency of couples with the female GG or GT genotypes (compared to the frequency of couples with the male *ZP2* GG or GT genotypes that are not expressed and should be a random draw from the allele frequencies in the population). However, we observed the same number of GG+GT genotypes (31 out of 37 total) among females (predicted to have lower fertility) and among males (not predicted to affect fertility).

Similarly, in our previous study we observed a weak but significant association between the paternal *ZP3R* C allele (rs4844573) and lower birth rate (Hart et al., 2018). However, we observed a similar number of couples with a male CC or CT genotype (21) who might have experienced infertility associated with that male allele compared to couples with a female CC or CT genotype (24) whose infertility could not be associated with that female allele. Both of those comparisons to fertility variation in Hutterites suggest that in our sample of couples seeking fertility assistance at PCRM the role of genotype variation at these three SNPs must be small.

A second and more important explanation for our negative results comes from other recent genomic analyses that identified an error in our previous understanding of human *ZP3R* and the identity and function of variation at the human SNP rs4844573. The human orthologs of mouse *Zp2* and *Zp3* are well known (Buffone et al., 2008), but until recently the identity of human *ZP3R* was not fully understood. Rohlfs et al. (2010) analyzed SNP variation in a protein-coding sequence on human chromosome 1 (chr1) that they called *ZP3R*, including the balanced polymorphism at the human SNP rs4844573. It is now clear that this gene identification was an error caused in part by the evolutionary history of gene duplications and pseudogenization events that are known to have occurred in both the mouse and human lineages.

Several genes in one region of mouse chr1 include *Zp3r* and other members of the C4b-binding protein family of genes and pseudogenes that each encode a series of CCP (or sushi) domains (McLure et al., 2004). Mouse C4bp and human C4BPA are orthologous proteins that can form oligomers when expressed in blood plasma and moderate the inflammatory response to infection as one of the regulators of complement activation (RCA) in the innate immune system (Arenzana et al., 1995). *Zp3r* in the mouse genome is descended from *C4bp* by an ancient gene duplication event, and the two genes are adjacent to each other in the RCA cluster on mouse chr1, but *Zp3r* has acquired a novel expression pattern (in the testis and acrosome) and a novel function (in sperm binding to the egg coat).

By contrast, the specific region of the human RCA cluster that is syntenic with mouse *Zp3r* and *C4bp* includes only one functional gene (*C4BPA*) and a non-functional pseudogene (*C4BPAP1*) previously annotated as a paralog of *C4BPA*. One of those two loci is probably the human ortholog of mouse *Zp3r*. Rohlfs et al. (2010) showed that polymorphisms in the gene they called *ZP3R* were in linkage disequilibrium (LD) with polymorphisms in human *ZP3* (on chr7). They argued that functional interactions between those polymorphisms when expressed in the sperm acrosome and in the egg coat might respond to sexual selection for pairs of advantageous alleles (and lead to the maintenance of LD between those physically unlinked loci).

Hart et al. (2018) pursued that discovery by documenting evidence for selection on three balanced polymorphisms in the coding sequences of *ZP3R, ZP3*, and *ZP2*; LD between the two balanced polymorphisms in *ZP3R* and *ZP2* (like the LD between *ZP3R* and *ZP3* found by Rohlfs et al., 2010); and effects of maternal *ZP2* genotype and paternal *ZP3R* genotype on two measures of fertility (Kosova et al., 2012). Hart et al. (2018) and Morgan and Hart (2019) argued that the gene called *C4BPA* is the human ortholog of mouse *Zp3r* and we ignored the human pseudogene *C4BPAP1*.

A more recent and more careful analysis by Carlisle et al. (2024) of synteny between genes in the mouse and human RCA clusters, and of phylogenetic relationships among the members of the C4b binding protein gene family, showed that the human pseudogene called *C4BPAP1* is the human ortholog of mouse *Zp3r*. Carlisle et al. (2024) noted that *C4BPAP1* also shows highest RNA expression in the testis (Carithers & Moore, 2015), in contrast to the lack of testis expression for human *C4BPA*. Those expression data (which were also not accounted for in previous studies; Hart et al., 2018; Morgan & Hart, 2019) are consistent with a former function for human *C4BPAP1* like the function of mouse *ZP3R* in fertilization, and with the idea that the human genome formerly included a functional *ZP3R* gene that evolved into a nonfunctional pseudogene after archaic humans diverged from the common ancestor of chimpanzees and bonobos.

Together those results show unambiguously that human *ZP3R* (annotated in the human reference genome GRCh38/hg38 as the pseudogene *C4BPAP1*) is the functional homolog (and the genetic ortholog) of mouse *Zp3r*, but no longer encodes a functional protein expressed in sperm and no longer has epistatic interactions with egg coat proteins during fertilization. The most important implication of that discovery for our new analysis is that the human SNP rs4844573 is not a polymorphism in the human *ZP3R* gene that might affect fertility, and instead is conclusively shown to be in the human *C4BPA* gene, which is expressed in the innate immune response to infection and has no role in human fertilization. Understanding that error in naming and annotating those human genes in comparison to the mouse genes (for which there are good experimental studies of gene expression and function) helps to explain why our new genotyping analysis did not show a link between genotype combinations and infertility: the SNP we analyzed is not expressed in sperm and could not directly affect fertility through gamete incompatibility.

## Conclusion & recommendations

The gene annotation error in Rohlfs et al. (2010) affected our previous work (Hart et al., 2018) and led directly to the error in designing this genotyping study. We focused on the SNP rs4844573 because it was in LD with SNPs in *ZP2* and in *ZP3* in previous studies and showed other evidence of evolving under selection with rs2075520 and with rs2906999. In light of the correct gene annotations by Carlisle et al. (2024) and the discovery that rs4844573 is not in a sperm-expressed gene, the genotype combinations observed among 37 couples in the experimental arm of our new study should not come as a surprise: those genotype combinations are not very different from random expectation (based on allele frequencies) because rs4844573 is not expressed in sperm and can’t interact with the other SNPs we analyzed in egg coat genes. For those reasons we stopped recruitment of participants into our study shortly after we learned about the correct gene annotations, but we completed our genotyping of the collected samples in order to conclude the study and document its negative results.

Although the coding sequence that includes rs4844573 is not expressed in sperm and cannot affect fertility, that SNP is associated with fertility variation (Hart et al., 2018) and is in LD with SNPs in both *ZP2* and *ZP3* (Rohlfs et al., 2010). Those patterns point to some unexplained functional or genetic association between genes in the RCA region of human chr1 and human fertility mediated by genes expressed in the egg coat or other cells. We currently lack a good hypothesis for those associations.

